# Parkinson’s-linked LRRK2-G2019S derails AMPAR trafficking, mobility and composition in striatum with cell-type and subunit specificity

**DOI:** 10.1101/2023.10.13.562231

**Authors:** Swati Gupta, Christopher A. Guevara, Alexander Tielemans, George W. Huntley, Deanna L. Benson

## Abstract

Parkinson’s (PD) is a multi-factorial disease that affects multiple brain systems and circuits. While defined by motor symptoms caused by degeneration of brainstem dopamine neurons, debilitating non-motor abnormalities in fronto-striatal based cognitive function are common, appear early and are initially independent of dopamine. Young adult mice expressing the PD-associated G2019S missense mutation in *Lrrk2* also exhibit deficits in fronto-striatal-based cognitive tasks. In mice and humans, cognitive functions require dynamic adjustments in glutamatergic synapse strength through cell-surface trafficking of AMPA-type glutamate receptors (AMPARs), but it is unknown how LRRK2 mutation impacts dynamic features of AMPAR trafficking in striatal projection neurons (SPNs). Here, we used *Lrrk2*^G2019S^ knockin mice to show that surface AMPAR subunit stoichiometry is altered biochemically and functionally in mutant SPNs to favor incorporation of GluA1 over GluA2. GluA1-containing AMPARs were resistant to internalization from the cell surface, leaving an excessive accumulation of GluA1 on the surface within and outside synapses. This negatively impacted trafficking dynamics that normally support synapse strengthening, as GluA1-containing AMPARs failed to increase at synapses in response to a potentiating stimulus and showed significantly reduced surface mobility. Surface GluA2-containing AMPARs were expressed at normal levels in synapses, indicating subunit-selective impairment. Abnormal surface accumulation of GluA1 was independent of PKA activity and was limited to D_1_R SPNs. Since LRRK2 mutation is thought to be part of a common PD pathogenic pathway, our data suggest that sustained, striatal cell-type specific changes in AMPAR composition and trafficking contribute to cognitive or other impairments associated with PD.

**SIGNIFICANCE STATEMENT:** Mutations in LRRK2 are common genetic risks for PD. *Lrrk2*^G2019S^ mice fail to exhibit long-term potentiation at corticostriatal synapses and show significant deficits in frontal-striatal based cognitive tasks. While LRRK2 has been implicated generally in protein trafficking, whether G2019S derails AMPAR trafficking at synapses on striatal neurons (SPNs) is unknown. We show that surface GluA1-AMPARs fail to internalize and instead accumulate excessively within and outside synapses. This effect is selective to D_1_R SPNs and negatively impacts synapse strengthening as GluA1-AMPARs fail to increase at the surface in response to potentiation and show limited surface mobility. Thus, LRRK2-G2019S narrows the effective range of plasticity mechanisms, supporting the idea that cognitive symptoms reflect an imbalance in AMPAR trafficking mechanisms within cell-type specific projections.

## INTRODUCTION

Leucine-rich repeat kinase 2 (LRRK2), a multi-functional kinase, has been the subject of intense study since it was discovered that inherited, autosomal dominant mutations that increase its kinase activity also increase the risk for Parkinson’s (PD). Elevated LRRK2 levels or kinase activity, in the absence of mutation, also occur in patients with idiopathic PD (1, 2), suggesting that the enzyme is part of a common disease pathology. Fronto-striatal based cognitive symptoms and altered corticostriatal processing can appear early in PD, prior to motor symptoms that are caused by the loss of dopamine neurons in the substantia nigra (3). Mechanisms driving such early symptoms and signatures are unknown, and there are no effective therapies to halt or reverse cognitive symptoms of PD.

LRRK2 expression levels are very low in dopamine neurons but enriched in striatal spiny projection neurons (SPNs). SPNs are obligatory processing units within looped brain circuits important for initiating or terminating action sequences that underlie goal-directed and habitual responses. SPNs receive the vast majority of their input from neocortical glutamatergic pyramidal cells and their outflow takes a direct or indirect path to the substantia nigra based on cellular identity defined principally by expression of either *Drd1* (encoding dopamine receptor D_1_R, direct-pathway SPNs) or *Drd2* (encoding D_2_R, indirect-pathway SPNs) (4). Lasting bidirectional changes in strength of glutamatergic synapses onto both SPN subtypes are thought to encode striatal-based learning and consistent with this idea, glutamatergic synapses in D_1_R and D_2_R SPNs undergo persistent strengthening (LTP) or weakening (LTD) (5–8). Previous work shows that in mice expressing a PD-associated *Lrrk2*^G2019S^ mutation, LTP is abolished in both subtypes, abrogating normal bidirectional synaptic plasticity required for striatal function (9, 10). As may be predicted by this more restricted synapse plasticity range, *Lrrk2*^G2019S^ mice exhibit significant dysfunction in striatally-based goal-directed and visuospatial attention tasks (11) that bear similarities to cognitive domains impaired in Parkinson’s (3, 12–14).

Based in part on work in other cell types, both the impairment in cognitive tasks that require striatal synapse plasticity coupled with abnormal synaptic plasticity in *Lrrk2*^G2019S^ SPNs has raised the possibility of defects in cellular mechanisms that target, support and regulate AMPA-type glutamate receptor (AMPAR) trafficking dynamics at striatal glutamatergic synapses (9, 15). Here, we test this directly. The data show that there is a SPN subtype- and AMPAR subunit-specific impact of *Lrrk2*^G2019S^ on AMPAR trafficking that serves to increase receptor stability on the surface at synaptic and non-synaptic sites. This has a profound impact on AMPAR subunit composition. Since AMPARs mediate most excitatory activity in the brain, the data suggest that a sustained increase in LRRK2 kinase activity broadly impacts information flow relevant to depression, anxiety and cognitive symptoms of early PD.

## RESULTS

### Surface pool of GluA1-containing AMPARs is increased in *Lrrk2*^G2019S^ SPNs

Previous work has shown that AMPAR-mediated activity is decreased in *Lrrk2* knockout SPNs while amplitude of AMPAR-mediated currents is increased in *Lrrk2*^G2019S^ SPNs during the first weeks of postnatal development (16–18). Based on these findings, we first asked whether AMPAR synthesis in striatal cells was altered in mice carrying a knockin mutation of *Lrrk2*^G2019S^ (16). We used bulk RNAseq to compare expression of AMPAR subunit mRNAs in *Lrrk2*^G2019S^ and wildtype mouse striata at P21. A volcano plot revealed that the mutation had a limited impact on transcription overall (**Suppl. Fig. 1A**) and there were no significant differences in levels of mRNAs encoding AMPAR subunits GluA1-4 (**Suppl. Fig. 1B**). Comparisons of AMPAR subunit transcript levels between patient-derived *LRRK2*^G2019S^ iPSCs and isogenic controls (GSE183499) and between putamen isolated from human PD patients and age-matched controls (GSE136666) (19) also showed no *LRRK2*^G2019S^-dependent or PD-associated differences in levels of AMPAR subunit mRNAs (**Suppl. Fig. 1B**). The absence of an effect of *Lrrk2*^G2019S^ on AMPAR transcription in striatum is also consistent with previous data showing that total levels of GluA1 and GluA2 proteins are similar in striatal lysates from *Lrrk2*^G2019S^ and wildtype mice (9, 15).

**Figure 1.**
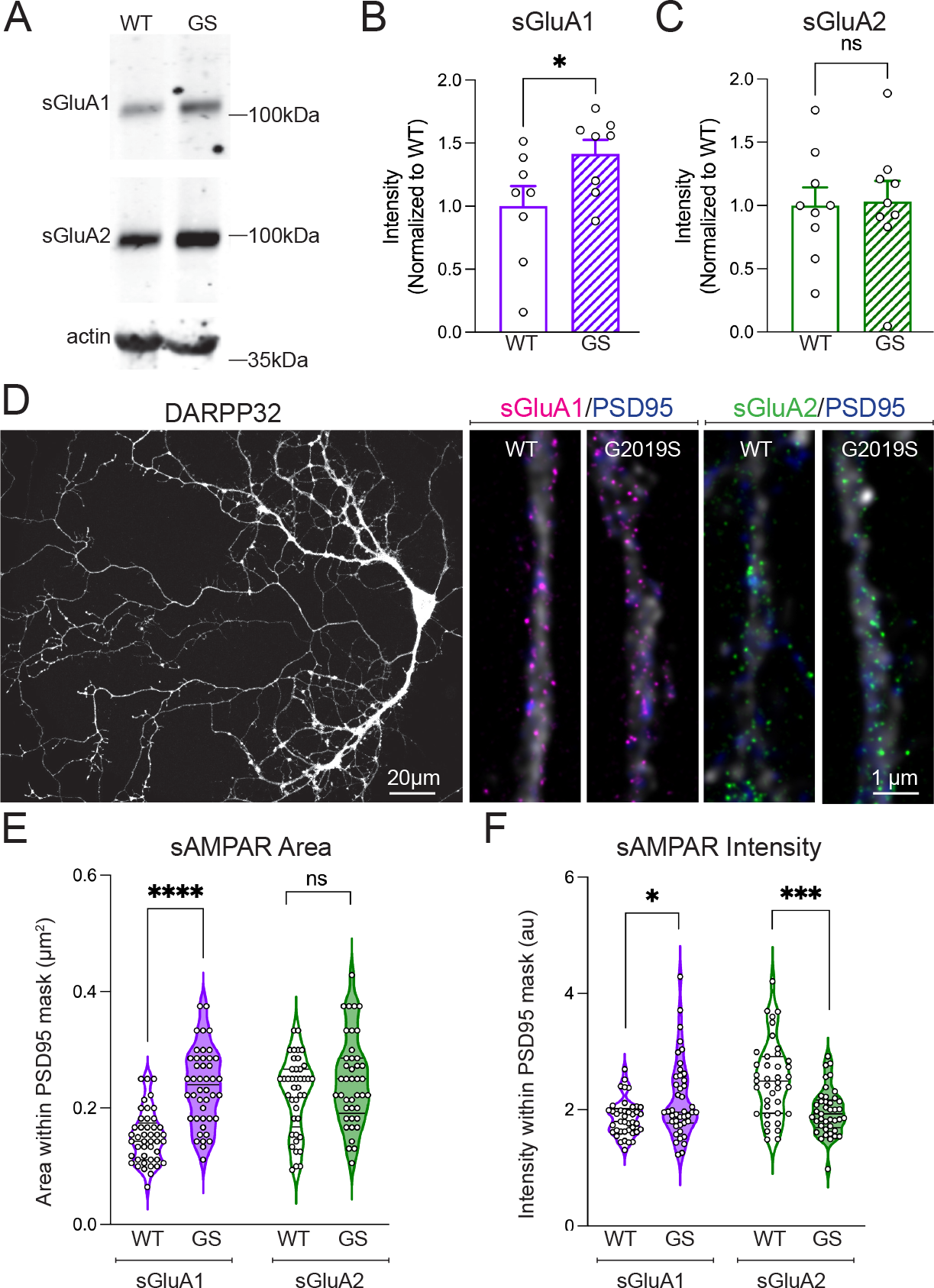
Lrrk2^G2019S^ disrupts AMPAR stoichiometry and subsynaptic distribution in SPNs. (**A - C**) Surface biotinylation was used to isolate endogenous surface (s) GluA1 and sGluA2 in acute striatal slices derived from wildtype (WT) and *Lrrk2*^G2019S^ (GS) mice. **A**) Representative Western blot images of sGluA1, sGluA2, and actin from dataset quantified in **B** and **C**. Scatterplot/bar graphs plot values normalized to WT ± SEM (n = 7 - 8 mice/genotype, 3 slices/mouse). Unpaired t test *p= 0.0492, compared to WT. **D**) Confocal image of DARPP32-labeled (white) co-cultured SPN (left) and super-resolution, STED images (right) of dendritic processes. Punctate sGluA1 (magenta), sGluA2 (green) labeling (tagged prior to permeabilization) associates largely, but not completed with PSD95 labeling (blue). **E** and **F**) Violin plots compare surface AMPAR area (**E**) and intensity (**F**) within masks defined by PSD95 labeling in SPNs. Unpaired t test, ****p<0.0001; ***p<0.0001; Mann Whitney test *p=0.04. n= 3 preps and 15 ROIs/genotype.

We next asked whether *Lrrk2*^G2019S^ regulates AMPAR subunit distribution between internal (i) and cell surface (s) receptor pools. To test this, we covalently tagged all cell-surface proteins with a reactive biotin-ester in acute striatal slices from *Lrrk2*^G2019S^ and wildtype mice, extracted the biotinylated proteins from lysates using streptavidin coated beads and identified AMPAR subunits by western blotting. These data indicate that sGluA1 levels, but not sGluA2 levels, were elevated significantly in *Lrrk2*^G2019S^ striatum compared to wildtype (**Fig. 1A-C**).

Since GluA1 is also expressed by striatal astrocytes and interneurons in addition to SPNs, we examined whether the enhanced sGluA1 seen in whole striatum reflected postsynaptic sites in SPNs. For this we used τ-STED (τ-<u>s</u>timulation <u>e</u>mission <u>d</u>epletion <u>m</u>icroscopy), a super-resolution approach (20), to quantify sGluA subunits (labeled prior to permeabilization) within a region defined by the postsynaptic density marker PSD95 in dissociated corticostriatal co-cultures. SPNs were identified and segmented using labeling for DARPP-32, a cytoplasmic protein that fills SPNs (**Fig. 1D-F**) but not other striatal cell types. Total area and intensity of sGluA1 clusters within PSDs were significantly greater in *Lrrk2*^G2019S^ SPN postsynapses compared to wildtype. In contrast, total area of sGluA2 clusters within PSDs was similar between genotypes, but sGluA2 intensity was diminished at *Lrrk2*^G2019S^ PSDs compared to wildtype, suggesting a modest reduction in sGluA2-containing AMPARs at postsynapses (**Fig. 1C-F**).

### Endocytosis of GluA1-containing AMPARs is greatly impeded in *Lrrk2*^*G2019S*^ SPNs

Increased levels of sGluA1-containing AMPARs could reflect impaired constitutive endocytosis. To probe this, we tracked and compared the extent of endogenous sGluA1 internalization in *Lrrk2*^G2019S^ and wildtype SPNs using an antibody feeding assay (**Fig. 2A**). A direct-conjugated, ATTO 594-tagged GluA1 rabbit polyclonal antibody (red) was used to label sGluA1 in SPNs that were cooled to prevent endocytosis and then fixed (t_0_) or chased at 37°C for 60 mins (t_60_), when AMPAR subunit internalization has plateaued (21, 22). All tagged sGluA1 was then labeled with anti-rabbit Alexa 647 (green) after which neurons were mildly permeabilized to label for DARPP-32 (**Fig. 2B, C)**. In wildtype SPNs at t_0_, intensity profiles along lines drawn through individual clusters show nearly complete overlap between red and green surface signals but by t_60_, are dominated by internalized (red-only) signal (**Fig. 2D vs. E**), as expected. In contrast, intensity profiles in *Lrrk2*^G2019S^ dendrites appear similar at t_0_ and t_60_ (**Fig. 2F vs. G**), suggesting sGluA1 failed to internalize over time. To compare internalization across time and genotype, we measured intensity of both fluorophores and generated an internalization index defined as the percent change in the ratio of surface/total GluA1 intensity **(Fig. 2H**). At t_0_ the mean internalization index value is near zero for both genotypes, but the value declines at t_60_ as expected in wildtype SPNs. In contrast, the internalization index remains virtually unchanged in *Lrrk2*^G2019S^ SPNs between t_0_ and t_60_. These data show that in *Lrrk2*^G2019S^ SPNs, GluA1 containing AMPARs failed to internalize. Since AMPARs are internalized by clathrin-mediated endocytosis (CME) (23–25), we tested whether CME was generally deficient in *Lrrk2*^G2019S^ SPNs by assaying transferrin receptor internalization (a marker for this pathway). However, transferrin receptors were internalized normally and similarly between genotypes (**Suppl. Fig. 2**), suggesting the impact of *Lrrk2*^G2019S^ on internalization of surface receptors is selective.

**Figure 2.**
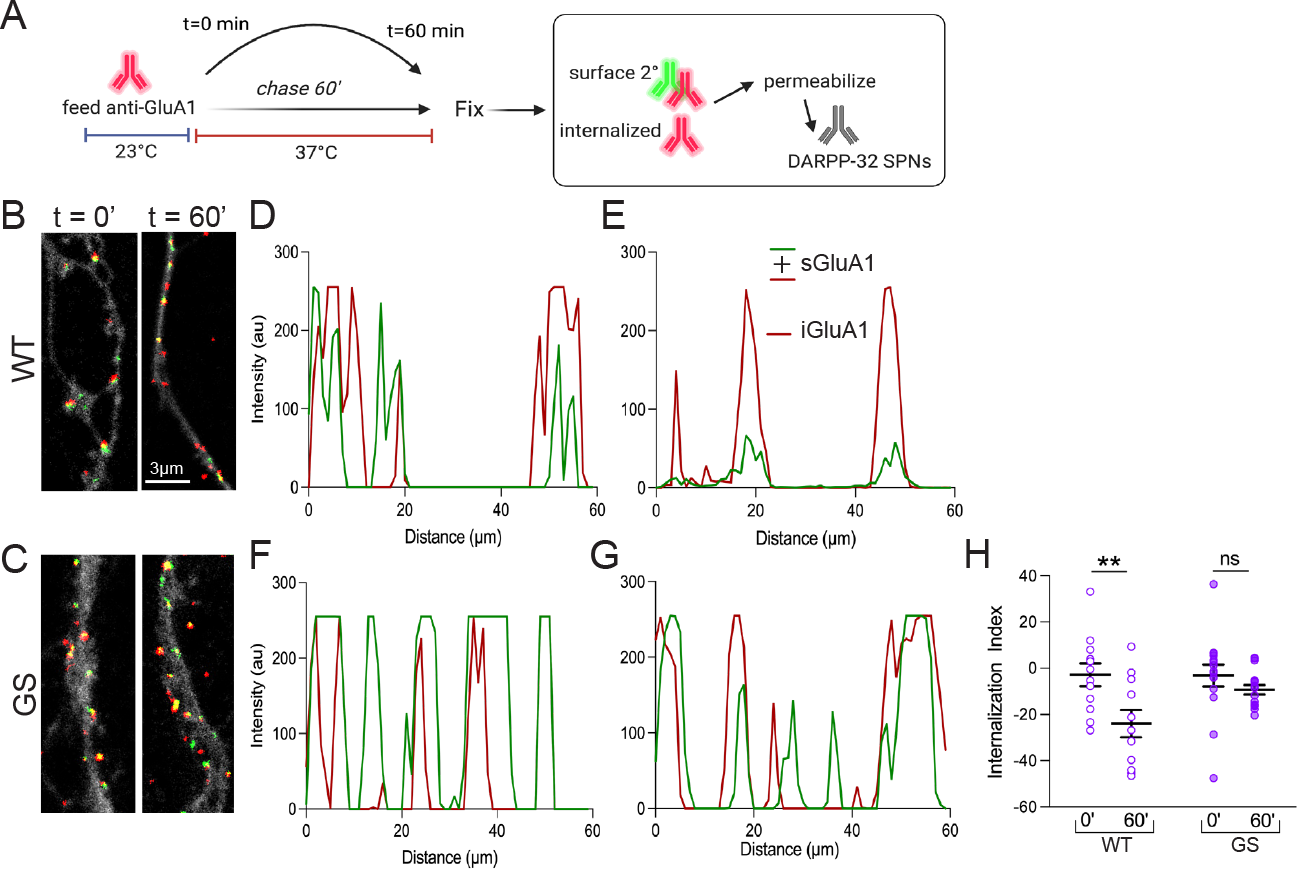
*Lrrk2*^G2019S^ disrupts GluA1 internalization in SPNs. **A**) Schematic outlines antibody feeding assay used to monitor GluA1 internalization in wildtype (WT) and *Lrrk2*^G2019S^ (GS) corticostriatal co-cultures (DIV16-18) and serves as a key for the colors used to show data. **B and C**) Overlay images show labeled surface (s) GluA1 (green mask + red mask) and internalized (i) GluA1 (red mask only) signal contained within DARPP-32 labeled SPNs (shown at a reduced intensity to permit visualization of puncta. Masks were generated in Image J and magnification is shown in B. **D - G**) Intensity distribution of green and red labeling along a 60 µm line scan. **H**) Quantification of the internalization index of GluA1 receptors in WT and GS SPNs at 0 and 60 min (n = 12 - 16 cells, 3 preps/genotype). Two-way ANOVA (F (1, 50) = 9.809, p = 0.0029), post hoc Šidák test **p = 0.0049.

### An LTP stimulus fails to increase sGluA1-containing AMPARs at *Lrrk2*^G2019S^ SPN synapses despite intact PKA-mediated GluA1-S845 phosphorylation

LTP of corticostriatal synaptic strength in SPNs may involve rapid insertion of calcium-permeable (CP) AMPAR subunits such as GluA1 (26), similar to hippocampal CA1 pyramidal neurons (27). The elevated accumulation of surface GluA1-containing AMPARs in *Lrrk2*^G2019S^ SPNs suggests that the inability to express corticostriatal LTP (9) may reflect GluA1 saturation, preventing the recruitment or insertion of additional GluA1-containing AMPARs needed to increase synapse strength. To examine this, we asked whether we could drive an increase in surface AMPAR subunits in *Lrrk2*^G2019S^ SPNs using a chemical-LTP (cLTP) stimulus applied to corticostriatal co-cultures (28). LTP in SPNs is mechanistically similar to PKA-dependent LTP in hippocampus in which CP-AMPARs (those lacking GluA2) are newly recruited to postsynaptic membranes within five mins following induction (6, 27, 29, 30). Five minutes following cLTP stimulation, cell surface proteins were biotinylated, isolated, and probed for GluA1 and GluA2 by western blot (**Fig. 3A, C**). In wildtype neurons, cLTP increased sGluA1 levels significantly compared to ACSF controls and the effect was blocked by the NMDAR inhibitor AP5, as expected (**Fig. 3A-C**). In contrast, *Lrrk2*^G2019S^ neurons failed to further increase sGluA1 levels in response to cLTP treatment compared to ACSF-treatment, and exposure to AP5 had no significant effect on sGluA1 levels compared to ACSF controls (**Fig. 3B, C**). Additionally, sGluA1 levels were elevated in ACSF control *Lrrk2*^G2019S^ neurons compared to wildtype neurons (**Fig. 3C**), as expected (**Fig. 1**). There were no differences in sGluA2 levels across treatments or genotypes (**Fig. 3C, D**). To confirm that the data from mixed cultures reflect SPNs, we measured and compared surface immunolabeling for GluA1 and GluA2 in DARPP-32-identified wildtype and *Lrrk2*^G2019S^ SPNs (**Fig. 3E - H**). These data matched the biochemical findings: there was a significant, NMDAR-dependent increase in sGluA1 levels in wildtype SPNs 5 minutes after cLTP, but this effect was absent in *Lrrk2*^G2019S^ SPNs (**Fig. 3E, G**), while sGluA2 levels were unchanged across genotypes or treatment conditions (**Fig. 3F, H**).

**Figure 3.**
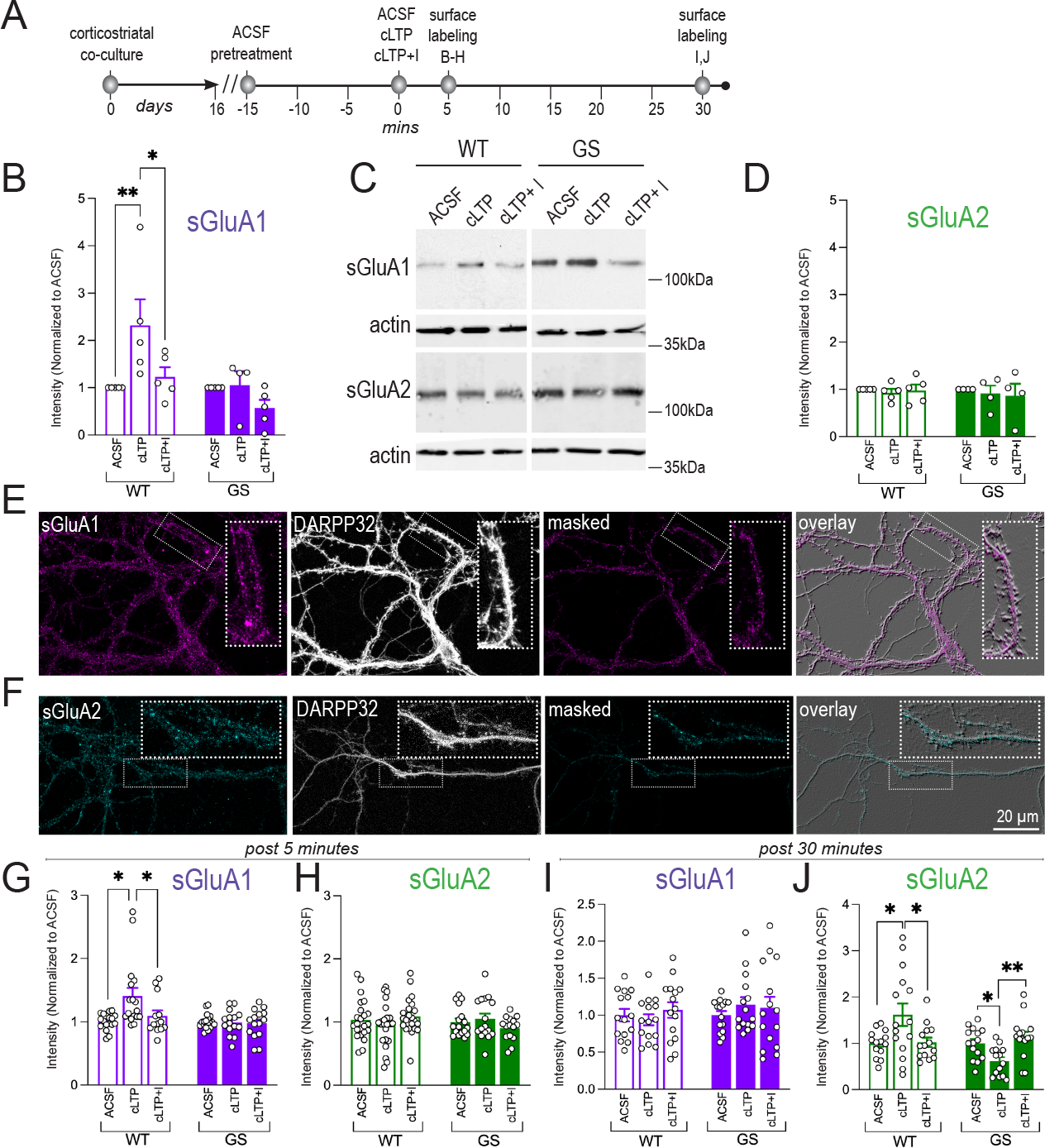
Surface GluA1-AMPARs fail to increase in response to cLTP in Lrrk2^G2019S^ SPNs. **A**) Schematic depicting experimental design. **A - C**: Representative western blot data (**C**) and scatterplot/bar graph showing quantification of surface sGluA1 (**B**) and sGluA2 (**D**) in WT and GS co-cultures in response to cLTP or cLTP + I (NMDA antagonist, AP5) and compared to ACSF controls ± SEM. One way ANOVA, (F(5, 23) = 4.661, p = 0.0044), post hoc Šidák’s test: **p = 0.0097, *p=0.039. (n = 4 - 5 preps/genotype). **E - F**: Representative confocal images of co-cultured WT and GS SPNs, immunolabeled for DARPP-32 (white, **E, F**), sGluA1 (magenta, **E**) or sGluA2 (cyan, **F**), following treatment with ACSF, cLTP or cLTP+I. **G - J**: Scatterplot/bar graphs show intensity levels (normalized to ACSF control for each group ± SEM) of sGluA1 and sGluA2 at 5 (**G, H**) and 30 (**I, J**) min post treatment (**A**). (**G**) WT: Kruskal-Wallis test **p = 0.007; post hoc Dunn’s multiple comparison test, ACSF vs cLTP *p = 0.0121, cLTP vs. cLTP+I *p = 0.0399; (**G, H**) (n= 11-17 cells, 3 preps/treatment). (**J**) WT: One-way ANOVA (F (2, 40) = 4.787, *p=0.0137), post hoc Tukey’s multiple comparison test ACSF vs cLTP *p = 0.0263, cLTP vs. cLTP+I *p = 0.0325. GS: One-way ANOVA (F (2, 41) = 7.618, **p=0.0015), post hoc Tukey’s multiple comparison test, ACSF vs cLTP *p = 0.0208, cLTP vs. cLTP+I **p = 0.0014 (I, J) (n = 14-15 cells, 3 preps/treatment).

Following LTP induction, the GluA2-lacking AMPARs are replaced by GluA2-containing AMPARs after 30 minutes (27, 31). Thus, to determine whether *Lrrk2*^G2019S^ also affected activity-dependent trafficking of GluA2-containing AMPARs, we compared sGluA2 levels in wildtype and *Lrrk2*^G2019S^ SPNs 30 min following cLTP stimulation (**Fig. 3A**). In wildtype SPNs, sGluA2 levels were increased significantly at 30 min compared to ACSF controls, while GluA1 levels had returned to control values, consistent with a receptor swap as expected. Increased sGluA2 levels 30 min post-cLTP were blocked by AP5 treatment (**Fig. 3I**). In contrast, *Lrrk2*^G2019S^ SPNs showed a significant *decrease* in sGluA2 levels 30 mins following cLTP—an effect that was blocked by AP5 (**Fig. 3J**). Additionally, at 30 mins, sGluA1 levels in the mutant SPNs remained elevated; there were no differences in sGluA1 levels between 5 and 30 min post-cLTP treatment (**Fig. 3I**).

PKA-mediated GluA1-S845 phosphorylation (pGluA1-S845) can drive AMPAR subunit insertion into the plasma membrane of SPNs (32–35), although there is debate whether this is required for LTP (36). While prior work has shown that LRRK2 can bind to and regulate the localization of PKARIIß, which negatively regulates PKA activity (37, 38), the G2019S mutation does not appear to impact this interaction (15, 37). Nevertheless, given the potential relevance of PKA activity here, we tested whether PKA-mediated pGluA1-S845 was disrupted in mutant SPNs by treating acute striatal slices from wildtype and *Lrrk2*^G2019S^ mice with forskolin, a potent activator of adenyl cyclase that increases PKA activity, followed by surface biotinylation and western blotting to compare total and surface levels of pGluA1-S845. In untreated striatum, levels of pGluA1-S845 phosphorylation were negligible, with no significant differences observed between genotypes (39) (**Suppl. Fig. 3A, B**). Forskolin treatment significantly and similarly increased pGluA1-S845 levels measured in the input (cytosolic) fractions in both genotypes (**Suppl. Fig. 3A, C**). Forskolin treatment also increased pGluA1-S845 significantly in wildtype striatum, as expected (**Suppl. Fig. 3A, B**). However, forskolin treatment of *Lrrk2*^G2019S^ striatum failed to drive a significant increase in surface pGluA1-S845 (**Suppl. Fig. 3A, B**). These data support that PKA activation and phosphorylation of GluA1 are intact in *Lrrk2*^G2019S^ striatum, but insertion, retention, or maintained pGluA1-S845-AMPARs in the plasmalemma is compromised.

### Enriched and Highly Stabilized Pool of Synaptic CP-AMPARs in Lrrk2^G2019S^ D_1_R SPNs

D_1_R and D_2_R SPNs drive competing circuits mediating movement, reinforcement learning, and reward. To test whether the excessive accumulation of sGluA1 in *Lrrk2*^G2019S^ SPNs is functional and to evaluate whether such impaired GluA1 trafficking dynamics showed D_1_R or D_2_R SPN subtype specificity, we crossed *Lrrk2*^G2019S^ mice with a fluorescent reporter line expressing *Drd1*a-tdTomato (40). We used whole-cell patch clamp recordings in acute striatal slices to compare the sensitivity of cortically evoked AMPAR-mediated postsynaptic currents (eEPSCs) to NASPM, a selective blocker of GluA2-lacking, CP-AMPARs (41). In wildtype D_1_R SPNs (tdTomato-positive), NASPM decreased eEPSC amplitudes by about 40% compared to eEPSC amplitudes prior to NASPM exposure. In contrast, we found that eEPSCs in *Lrrk2*^G2019S^ D_1_R SPNs were significantly more sensitive to NASPM, decreasing eEPSC amplitudes by about 70% (**Fig. 4A, B**), consistent with a significant enrichment of synaptic GluA1 shown biochemically and anatomically above. In contrast, NASPM had little effect on eEPSCs in wildtype or *Lrrk2*^G2019S^ putative D_2_R SPNs (tdTomato-negative, **Fig. 4B, C**). These data show that the enhanced sGluA1 levels observed biochemically and anatomically in *Lrrk2*^G2019S^ SPNs are functionally relevant, and that altered AMPAR subunit stoichiometry at corticostriatal synapses predominantly reflects D_1_R SPNs.

**Figure 4.**
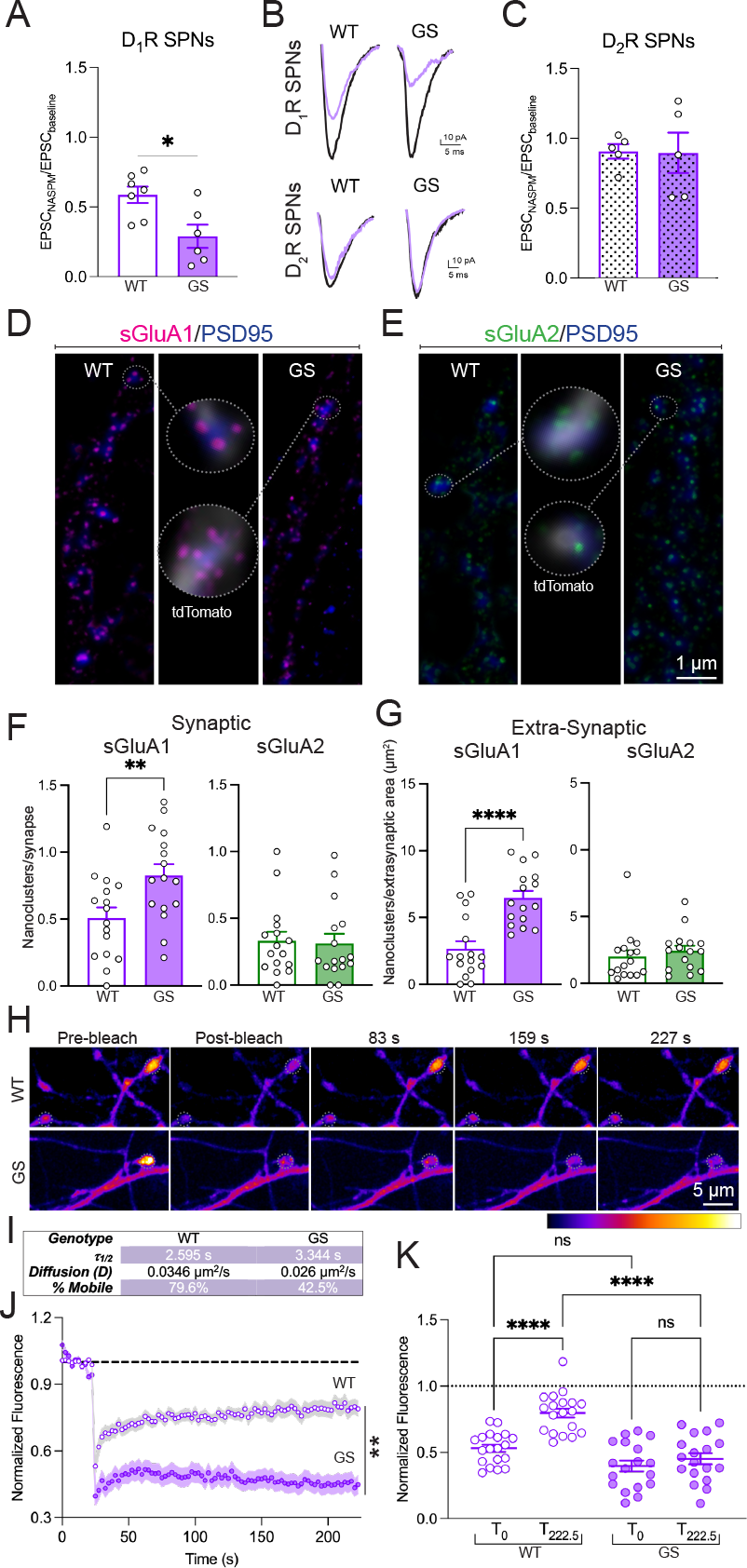
Lrrk2^G2019S^ D_1_R SPNs have an excess of stabilized CP-AMPARs. **A, C**) Bar graph/scatterplots comparing functional contribution of CP-AMPARs using ratios of EPSCs evoked in the presence/absence of NASPM in D_1_R (**A;** *Drd1tdTom+*) and D_2_R (**C;** *Drd1tdTom-*) SPNs in WT and GS mice (P70 - P90) in acute slices through dorsal striatum. Bars are mean ± SEM (n = 11 cells, 5 - 6 mice/group, unpaired t test *p = 0.0122). Example traces (**B**) show AMPAR currents before (black) and after (lavender) bath application of NASPM (200µM, 10 min). **D - G**, Superresolution (tau-STED) images (**D, E**) and quantification (**F, G**) of synaptic and extrasynaptic receptors in 21 DIV WT and GS D_1_R SPNs expressing tdTomato (white, *Drd1*Cre/+; *Ai14*, in D, E, and used to segment D1R SPNs for F, G) and co-cultured with unlabeled cortical neurons of the same genotype. sGluA1 (**D**, magenta, STED) and sGluA2 (**E**, green, STED) puncta in relation to PSD95 labeled postsynaptic sites (blue, confocal). Circled zones are enlarged in the center panels. **F, G**) Bar graph/scatterplots show synaptic (**F**, within a mask defined by PSD95) and extrasynaptic (**G**, outside a PSD95 mask) sGluA1 (lavender) and sGluA2 (green) nanoclusters in D_1_R SPNs. **F**: Unpaired t test **p = 0.0369; n = 16 ROIs/genotype). **G**: Unpaired t test ****p<0.0001; n = 16 ROIs/genotype. **H - K**) Examples (**H**) and quantification (**I - K**) of FRAP experiments. **(H)** Time lapse confocal images pre- and postphotobleaching (dotted circles approximate ROIs) in WT and GS D_1_R SPNs labeled and co-cultured as above. Table (**I**) compares time constant, diffusion (D) and % mobile receptors. D was calculated using: D = 0.25 (r^2^/τ_1/2_), where r refers to the bleach radius and τ_1/2_ to the time constant^46^. Graph (**J**) plots normalized SEP-GluA1 fluorescence recov-ery in WT and GS neurons imaged every 2.5 s. Lighter shading is ± SEM. Two-way RM-ANOVA (F(89, 3115) = 17.79, **p < 0.001, n = 19-20 spines/genotype. Scatterplot (**K**) compares recovery of SEP-GluA1 intensity at T222.5 s time point relative to post-bleach in WT and GS D_1_R SPNs. One-way ANOVA (F(3, 70) = 24.15, p < 0.0001), post hoc Tukey’s multiple comparison test ****p<0.0001.

Receptor subunit nanoclusters detected using STED can be used to estimate receptor composition (28). Thus, to compare AMPAR composition between genotypes in D_1_R SPNs, we used τ-STED to resolve and quantify numbers of sGluA1 and sGluA2-containing nanoclusters within PSDs in D_1_R SPNs cultured from wildtype or *Lrrk2*^G2019S^ mice that were identified by D_1_R-Cre-dependent expression of tdTomato. *Lrrk2*^G2019S^ D_1_R SPNs exhibited greater numbers of synaptic sGluA1 nanoclusters but no difference in synaptic sGluA2 nanoclusters compared to wildtype D_1_R SPNs (**Fig. 4D, E**), supporting that there is an altered stoichiometry of sAMPAR subunits at mutant D_1_R SPN synapses, with an increased proportion of GluA1 and a decreased proportion of GluA2. We next asked whether sGluA1-containing AMPARs were also accumulating at extrasynaptic sites in *Lrrk2*^G2019S^ D_1_R SPNs by counting nanoclusters on dendrites outside regions labeled by PSD95. Numbers of sGluA1-but not sGluA2-containing nanoclusters were also enhanced extrasynaptically in *Lrrk2*^G2019S^ D_1_R SPNs, suggesting that GluA1 exocytosis may be intact while deficits in GluA1 endocytosis may drive surface accumulation (**Fig. 2**). Since increased synaptic GluA1 is often positively correlated with synapse size, we also assessed density and size of PSD95-labeled clusters in D_1_R SPN dendrites, but no genotype-dependent differences were detected (**Suppl. Fig. 4**), consistent with the absence of an impact of *Lrrk2*^G2019S^ on spine size (15, 42).

An increase in extrasynaptic sGluA1-containing AMPARs might serve as a resource for synaptic trafficking when conditions demand (43, 44), but the failure to rapidly recruit GluA1-containing AMPARs in *Lrrk2*^G2019S^ SPNs in response to cLTP (**Fig. 3**), suggests this pool is unavailable. To compare the dynamics of sGluA1-AMPAR trafficking in D_1_R-SPNs, we measured fluorescence recovery after photobleaching (FRAP) of SEP-tagged GluA1 expressed in wildtype and *Lrrk2*^G2019S^ D_1_R SPNs expressing td-Tomato. SEP is a pH-sensitive GFP variant that emits fluorescence at the cell surface (pH 7.4), but is quenched in the acidic environment of intracellular vesicles (< pH 6; (45)). Re-population of a photobleached area with SEP-GluA1 receptors results from lateral diffusion from neighboring unbleached areas and exocytosis from intracellular compartments. We photobleached 1.2 µm diameter ROIs that were centered on the heads of dendritic spines using a 488nm laser, and fluorescence recovery was monitored every 2.5 s for four mins subsequently using acquisition conditions that showed stable SEP-GluA1 intensity in unbleached regions and tdTomato intensity (**Fig. 4H, Suppl. Fig. 5A-D**). The effectiveness of the photobleaching was similar between genotypes (**Fig. 4H, J, K, Suppl. Fig. 5A, B**). Plots of normalized SEP-GluA1 fluorescence over time showed that the wildtype recovery curve fit a nonlinear bi-phasic association, with a τ_fast_ = 3.74s and τ_slow_ = 93.63s, while the *Lrrk2*^G2019S^ recovery fit a nonlinear mono-phasic association with a single τ = 4.76s, closer to the τ_fast_ seen in wildtype (**Fig. 4J**). SEP-GluA1 recovered to only 45.2% of baseline in *Lrrk2*^G2019S^ D_1_R SPN spines in comparison to 79.6% recovery in wildtype D_1_R SPN spines (**Fig. 4I - K**). Based on the τ values, we extrapolated diffusion coefficients for the mobile fraction of receptors (46), which show that SEP-GluA1 AMPARs in wildtype D_1_R SPN spines diffuse at a rate 33% faster (0.035 μm^2^/s) than those in *Lrrk2*^G2019S^ D_1_R SPN spines (0.026 μm^2^/s) (**Fig. 4I**). Thus, the excessive surface GluA1-containing receptors in *Lrrk2*^G2019S^ D_1_R SPNs show greatly diminished mobility.

## DISCUSSION

Our data show that Parkinson’s associated *Lrrk2*^G2019S^ mediates a significant and selective impact on AMPAR composition in dorsal striatum that abnormally increases the surface synaptic and extrasynaptic incorporation of GluA1, but not GluA2, in D_1_R SPNs, but not in D_2_R SPNs. Increased sGluA1-containing AMPARs result in part from their severely impeded internalization that contributes to a saturating ceiling on synapse plasticity by preventing additional GluA1-AMPAR insertion or recruitment from extrasynaptic pools under activity-dependent conditions. The functional impact is observed in preparations from early postnatal and young adult mice. Together the data outline a powerful and selective impact on AMPAR trafficking, composition and mobility early in life in the presence of a pathological increase in LRRK2 kinase activity that may contribute to cognitive or other functions that normally rely on intact AMPAR trafficking dynamics, but deteriorate in Parkinson’s.

We found a potent impact of *Lrrk2*^G2019S^ on AMPAR recycling and dynamics by several biochemical and imaging approaches where excessive accumulation of sGluA1 was observed both within and outside of PSD95-defined postsynaptic sites. Our analyses further support that such an increase results from dysfunctional endocytosis, as endogenous GluA1 containing-AMPARs showed very little receptor internalization in mutant SPNs compared to wildtype. Excess sGluA1 levels could also result from more rapid receptor recycling, but this possibility is countered by the absence of SEP-GluA1 fluorescence recovery following photobleaching. The GluA1 internalization defect is likely selective in that transferrin receptor surface expression and internalization were similar between wildtype and mutant SPNs, consistent with previous work in iPSC-derived microglia expressing *LRRK2*^G2019S^ (47). Selectivity could be imparted by a targeted impact of LRRK2 on a clathrin-independent pathway (48) or on the juxta-synaptic endocytic zones that preferentially internalize AMPARs (49), and which display distinct clathrin dynamic properties (50).

The significantly greater sensitivity of evoked synaptic AMPAR-mediated responses to NASPM revealed by whole cell recordings under baseline conditions support that increased levels of synaptic GluA1-AMPARs likely correspond to CP-AMPARs comprising GluA1 homomers. Consistent with this enrichment, synaptic delivery of GluA1-AMPARs in response to cLTP (27, 51) failed in *Lrrk2*^G2019S^ SPNs, suggesting these synapses have hit a ceiling of saturation that cannot support further activity-driven insertion of additional GluA1 subunits. It may be that receptors are too firmly anchored or cannot easily transit to the PSD. Additionally, the excess extrasynaptic sGluA1-AMPARs in *Lrrk2*^G2019S^ SPNs do not appear to serve as a reserve pool for rapid synaptic deployment since our FRAP data, which showed significantly decreased recovery and diffusion rates for SEP-GluA1, strongly support that the immobile fraction of sGluA1-AMPARs is enhanced in *Lrrk2*^G2019S^ D_1_R SPNs. Alternatively (or additionally) exocytosis or trafficking to synapses may be negatively impacted. A subset of Rab-GTPases, including Rab8 and Rab10 which are typically associated with exocytosis, are validated substrates for LRRK2-mediated phosphorylation (52). Counter to expectation, but similar to what is observed here, hippocampal neurons expressing a dominant negative Rab8, show *increased* synaptic recruitment of exogenously expressed GluA1 and failed to produce LTP in response to potentiating stimuli (53). LRRK2-phosphorylated Rab8 and -10 can also sequester MyosinV proteins (54), which in neurons would be expected to impede the motor protein-dependent transport of AMPARs into spines during LTP (55). Although the basis for the SPN subtype-specific effects are unclear, it may be that relevant LRRK2-targeted effectors are differentially distributed between SPN subtypes.

Our data also suggest that *Lrrk2*^G2019S^ SPNs exert compensatory or adaptive responses to a synaptic excess in sGluA1. The increase in extrasynaptic sGluA1-AMPARs, which are neither readily internalized (**Fig. 2**) nor mobile (**Fig. 4**), may be actively excluded from synapses in order to maintain relatively normal synaptic transmission. Consistent with this idea, the excessive synaptic sGluA1 has a negligible impact on baseline EPSC amplitudes (15, 16, 18), spine size is unchanged (15, 42), and synaptic sGluA2 levels drop modestly (Fig. 1) in *Lrrk2*^G2019S^ SPNs. Altered responsiveness appears to emerge when the synapses are challenged, here by a strong potentiating stimulus, which revealed that sGluA1-AMPARs failed to be recruited during induction and sGluA2-AMPAR levels dropped as LTP normally consolidates. The drop in sGluA2-AMPARs may be indicative of a form of postsynaptic LTD that has been described in SPNs (56), and could be consistent with prior work in *Lrrk2*^G2019S^ striatum showing that a normally potentiating stimulus yields instead LTD (9).

## MATERIALS AND METHODS

### Mice

*Lrrk2*^*G2019S*^ knock-in mice were generated by Eli Lilly labs and characterized previously (16). Male and female homozygous *Lrrk2*^*G2019S*^ mice (C57BL/6-Lrrk2tm4.1Arte; RRID:IMSR_TAC:13940 (10–12 weeks old) and age- and strain-matched wildtype mice were used in all experiments unless specified otherwise. The *Lrrk2*^*G2019S*^ mice were backcrossed to wildtype C57BL/6NTac (RRID:IMSR_TAC:B6) every fifth generation to prevent genetic drift. For electrophysiology experiments male and female *Lrrk2*^G2019S+/-^ and wildtype mice heterozygous for *Drd1a* -tdTomato (B6.Cg-Tg(Drd1a-tdTomato)6Calak/J; RRID:IMSR_JAX:016204, The Jackson Laboratory (57)) were used. FRAP live imaging and super resolution STED imaging was performed in cultured D_1_R neurons derived from pups generated by crossing mice that express Cre recombinase under the control of the mouse Drd1a promoter (B6;129-Tg(Drd1-cre)120Mxu/Mmjax (RRID:MGI:5578135), The Jackson Laboratory) (58) with Ai14 mice (B6.Cg-Gt(ROSA)26Sortm14(CAG-tdTomato)Hze/J (RRID:IMSR_JAX:007914, The Jackson Laboratory) (59) that express robust tdTomato fluorescence following Cre-mediated recombination. Mice were genotyped using automated genotyping services offered by Transnetyx. All animals were kept with dams until weaning age (P21) and then housed in single-sex groups of 3 - 5 animals per cage. The care and treatment of all animals were in strict accordance with guidelines of the Institutional Animal Care and Use Committee of the ISMMS and those of the National Institutes of Health.

### Surface Biotinylation and Western Blotting Assays

Acute coronal brain slices (350 μM) were taken from isoflurane anesthetized male and female homozygous *Lrrk2*^G2019S^ mice (10–12 weeks old) and age- and strain matched wildtype (*Lrrk2*^WT^ or WT) adult mice. Following recovery of slices for 1 hour at 37°C in ACSF, 3 slices/mouse were transferred to ice-cold biotin solution (Thermo Scientific EZ Link Sulfo NHS SS Biotin at 0.6 mg/ml in ACSF) for 40 min, then quenched in 0.1mM Glycine in ACSF, 2 washes, 5 min, and washed in ACSF,5 min. Either whole striatum or dorsomedial striatum was subdissected as indicated in relevant figure legends. The tissue was solubilized for 1 h in RIPA buffer [50 mM Tris pH 7.5, 1 mM EDTA, 2 mM EGTA, 150 mM NaCl, 1% NP40, 0.5% DOC, 0.1% SDS, and protease (Thermo Fisher Scientific, Cat # A32965) and phosphatase (Thermo Fisher Scientific, Cat # A32957) inhibitor cocktails] and the lysates were then centrifuged to pellet cell debris. 10% of the supernatant was taken as a total protein sample and the remainder was incubated for 2h with 20μl immobilized NeutrAvidin beads (ThermoScientific, Cat # PI29200) 50% slurry at 4°C to precipitate biotin-labeled membrane proteins. Beads were washed two times in high-salt RIPA buffer (350mM NaCl) and a final wash in RIPA buffer and analyzed by SDS-PAGE and western blotting. Biotinylated surface proteins were identified by using antibodies, anti-GluA1 (1:500, NeuroMab clone N355/1, RRID: AB_2315839), anti-GluA2 (1:500, Millipore Sigma MABN1189, RRID:AB_2737079), pGluA1-S845 (1:500, Fisher Scientific AB5849, RRID:AB_92079) and anti-Actin (1:5000, Millipore Sigma MAB1501, RRID:AB_2223041) used as a loading control. Bands were detected using fluorophore coupled anti-mouse/rabbit secondary antibodies (DyLight 800, Cell Signaling Technology; RRID: AB_10697505; and DyLight 680, Pierce; RRID:AB_1965956) followed by imaging on LICOR Odyssey CLX imager (LI-COR Biosciences) or the Azure Imager. ImageJ (NIH) was used to quantify relative protein levels via band intensity normalized to a loading control (Actin).

### Culture Preparations

Cortical and striatal neurons were co-cultured from E16 -18 wildtype, *Lrrk2*^G2019S^, *Drd1aCre*^*+/-*^*;Ai14* and *Drd1aCre*^*+/-*^*;Ai14;Lrrk2*^G2019S+/-^ mice. Briefly neurons were dissociated and plated on18 mm coverslips coated with 1 mg/ml poly-L-lysine as described previously (60) and maintained in a humidified 37°C incubator with 5% CO_2_ until DIV16 - 22. Cortical and striatal neurons (300,000 cells/6 cm dish) were co-plated at a ratio of 3:2 to ensure normal morphological and physiological development and function of SPNs in culture (61). As Drd1a expressing neurons can also be found within the cortex, brains extracted from *Drd1aCre*^*+/-*^ *;Ai14* and *Drd1aCre*^*+/-*^*;Ai14;Lrrk2*^G2019S+/-^ pups were screened for tdTomato expression on an epifluorescent microscope and cortical neurons were extracted from tdTomato-negative brains while striatal neurons were extracted from tdTomato-positive brains to selectively culture labeled D_1_R SPNs. For transfection of pCAG-SEP-GluA1 cDNA (gift of R. Huganir, Johns Hopkins, MD), co-cultured neurons were transfected using Lipofectamine 2000 (Invitrogen) at DIV 11-12 according to the manufacturer’s recommendations 48h prior to imaging experiments.

### Immunostaining

Cultured neurons were fixed with 4% PFA and 4% sucrose in TBS, pH 7.3. Coverslips were washed three times for 10 min in TBS and blocked in 5% BSA for 1h at room temperature. After blocking, coverslips were incubated with primary antibodies targeting surface epitopes (GluA1;1:500, Alomone AGP-009, RRID:AB_2340961 or GluA2; 1:500, Alomone AGC-005-GP, RRID:AB_2756617) at 4°C overnight. Coverslips were washed three X 10 min, permeabilized with 0.5% Triton X-100 in blocking buffer for 15 min and incubated with primary antibodies against internal epitopes (DARPP32 (1:500, Cell Signaling Technology 2306, RRID:AB_823479) or PSD95 (1:500, Thermo Fisher Scientific MA1045, RRID:AB_325399) at 4°C overnight. Coverslips were washed three X 10 min in TBS and incubated with secondary antibodies (Alexa Fluor® 647-conjugated Anti-Guinea Pig IgG Antibody (1:500, Jackson ImmunoResearch 706-605-148, RRID:AB_2340476), Alexa Fluor® 594-conjugated Anti-Rabbit IgG Antibody (1:500, Jackson ImmunoResearch 711-585-152, RRID:AB_2340621), Anti-Mouse IgG (whole molecule)-ATTO 488 antibody (1:500, Sigma Aldrich 62197-1ML-F, RRID:AB_1137649) diluted in blocking buffer for 1 h at room temperature, shielded from light. After incubation was completed, three 10-min washes at room temperature were performed. Coverslips were mounted on Superfrost Plus slides using Mowiol, sealed using nail polish, and dried overnight in the dark.

### Antibody Feeding Assay

Cultured neurons were live-labeled with a direct-conjugated, ATTO-594 (red) rabbit anti-GluA1 antibody (1:100, Alomone AGC-004-AR, RRID:AB_2340944) for 15 min at room temperature. The coverslips were washed, and a subset of coverslips was fixed immediately (t_0_) or chased at 37°C for 60 minutes (t_60_) and then fixed. Under non-permeabilizing conditions, the ATTO labeled surface-GluA1 receptors were labeled with Alexa Fluor Plus 647 conjugated anti-Rabbit IgG Secondary Antibody (1:200, Thermo Scientific A32733, AB_2633282) for 1h at room temperature. Thereafter using the immunostaining protocol described above, DARPP32 immunolabel was used to positively identify SPNs.

### Transferrin Receptor Endocytosis Assay

Cultured neurons were either fixed immediately (t_0_) or treated with 30μg/ml of ligand transferrin (Jackson Immunoresearch, 015-600-0500) at 37°C for 5 minutes (t_5_) and then fixed. Under non-permeabilizing conditions, transferrin receptors were labeled using primary antibody (1:100, Developmental Studies Hybridoma Bank G1/221/12, RRID: AB_2201506) and Alexa Fluor® 647-conjugated Anti-Mouse IgG Antibody (1:500, Thermo Fisher Scientific AB-0662) using the immunostaining protocol described above. The DARPP32 immunolabel was used to positively identify SPNs.

### Confocal Microscopy

Single optical images of cultured neurons plated on coverslips were captured on a Leica SP8 STED 3X Falcon confocal microscope (Leica Microsystems) using a 63x/1.4 PlanApo oil immersion lens, frame size set to 1024X1024; 16bits per pixel. A thresholding function was applied to generate a mask of the DARPP32-positive or td-Tomato-filled SPN that was overlaid to quantify mean immunofluorescent GluA intensity within SPNs using NIH ImageJ. Postsynaptic (or extrasynaptic) areas were defined using a second mask based on thresholding applied to PSD95 labeling.

### STED

Images were captured on a Leica SP8 STED 3X Falcon microscope using 100x 1.4 STED-HC PlanApo oil immersion lens (Leica Microsystems), an optical zoom of 6, frame size was set to 1024X1024 for < 20 nm pixel size. 633 nm laser derived from 80MHz pulsed White Light Laser (Leica Microstystems) was used to excite Aberrior-Star-635P (1:500, Aberrior ST635P-1006-500UG, RRID:AB_2893230) labeled GluA subunit of AMPARs. The 775 nm pulsed laser (at 70% power) was used to reduce the point-spread function (PSF) for the fluorophore. Tau STED was used to filter pixels based on phaser plots. Emission was collected with a Hybrid Detector (HyD, Leica Microsystem; gain 100). Images were deconvolved using Huygens Professional 22.10 (SVI, Netherlands), Classic Maximum Likelihood Estimation. Microscope parameters were uploaded from image metadata and deconvolution configuration was set to auto.

### FRAP

Coverslips were mounted in an imaging chamber immersed in Neurobasal media (without phenol red) in which the neurons were maintained at 37°C with 5% CO_2_. Transfected SEP-GluA1 and tdTomato positive D_1_R SPNs were identified manually through the oculars on a Zeiss LSM980 Airyscan 2 (Carl Zeiss Microscopy) using the widefield setting and 555nm LED illumination. A single optical image was then taken using 561nm diode laser illumination 63X 1.3NA oil-immersion objective, a zoom 4.5X, 512 X 512 pixels, 16 bits per pixel and a pinhole = 74 µm. Spines were identified using circular ROIs of diameter 1.2 μm. Acquisition parameters were set using ZEN Blue software to excite using the 561nm diode laser (at 4% power, tdTomato) and 488 nm Argon laser (at 3% power, SEP-GluA1) every 2.5s, with the first ten images acquired to obtain baseline, followed by high laser power (488nm at 50% for 5 iterations with a scan speed of 7) to photobleach select ROIs. Images were then acquired every 2.5s after bleaching to monitor tdTomato fluorescence (photobleaching) and to capture the recovery of fluorescence, which reflects diffusion from unbleached zones and delivery of SEP-GluA1 to the plasma membrane. Non-photobleached ROIs served to control loss in fluorescence because of imaging. The data were exported to Excel and the photobleached ROIs were analyzed by normalizing to non-photobleached ROI control to correct for any loss in fluorescence due to imaging, followed by subsequent normalization to the pre-bleach baseline. The data were plotted using Prism (GraphPad) and analyzed using non-linear exponential fit. Diffusion coefficients were calculated as described by Kang and colleagues (46).

### Chemical LTP

We used a well-characterized pharmacological paradigm to induce NMDAR-dependent LTP with the NMDAR co-agonist glycine (28). Briefly, cultured neurons were treated with ACSF (in mM: 14 NaCl, 5KCl, 2 CaCl_2_, 1 MgCl_2_, 30 glucose and 10 HEPES, pH 7.4) containing 0.5 µM TTX, 1 µM strychnine and 20 µM bicuculline) for 15 min and then switched to ACSF (control) or ACSF-Mg^2+^-free extracellular solution containing glycine (200 µM) (cLTP) or cLTP solution containing NMDAR antagonist (50 µM D-APV; D-2-amino-5-phosphonovalerate) to block LTP (cLTP+I) for 5 min or 30 min. Cultured neurons were either biotinylated to assay surface protein levels or fixed and subject to immunostaining.

### Electrophysiology

Whole cell patch clamp recordings were conducted in acute striatal slices as described above for surface biotinylation above. Recordings from SPNs located in dorsomedial striatum were conducted similarly to (62). Following recovery, slices were transferred to a recording chamber at 31°C containing Gabazine (GBZ; 10 µM; Sigma SR-95531). SPNs were visualized on an upright epifluorescence microscope (BX50WI; Olympus) using a 40X water-immersion lens along with an IR-1000 infrared CCD monochrome video camera (DAGE MTI) and D_1_R expressing SPNs were identified using epifluroescent illumination. Whole cell recordings were performed utilizing a glass borosilicate pipette with resistance of 2 - 4MΩ and filled with the following (in mM): 124 K-gluconate, 10 HEPES, 10 phosphocreatine di(Tris), 0.2 EGTA, 4 Mg_2_ATP, 0.3 Na_2_GTP. Functional Ca^2+^-permeable AMPAR (CP-AMPAR) were assessed in DMS SPNs as described previously (9). To evoke corticostriatal responses, a stimulating electrode was positioned medially at the layer 6/white matter border. Evoked AMPAR responses were taken from SPNs at baseline and followed by a bath-application of the CP-AMPAR antagonist 1-naphthyl acetylspermine (NASPM; Tocris 2756, 200 μM in 1:100 DMSO). Cells were incubated in NASPM for 10 mins before taking evoked AMPAR responses. Nine evoked EPSCs were taken at baseline and then after NASPM application. Data was collected using Clampfit software (pCLAMP, Molecular Devices) and analyzed using Mini Analysis module.

### Data Analysis and Statistics

Statistical comparison of two groups was conducted using two-tailed Student’s t test, and for non-paramatric data, Mann-Whitney test. For comparison of more than two groups, Bartlett’s test for equal variance was performed; one-way ANOVAs were conducted to compare across groups with equal variances. To compare data from multiple groups over time, two-way ANOVA was performed, with either *Š*idák’s or Tukey’s multiple comparison tests. For nonparametric data in multiple group comparisons, rank transformation was conducted on data and analyzed through a mixed-effects analysis, followed by *Š*idák’s posttest. A p value ≤ 0.05 was considered significant. Analyses used GraphPad Software Prism 9. Data are presented as mean values ± SEM. Numbers and particular tests used are given in the figure legends.

## Supporting information

Supplementary Figures & References

## Acknowledgements

This work was supported by R01NS107512 and R21NS130841 from the National Institute of Neurological Disorders and Stroke, the Michael J Fox Foundation MJFF 022867, and T32-MH087004 from the National Institute of Mental Health. Special thanks to Dr. Richard L. Huganir for providing the SEP-GluA1 construct, Dr. Abhishek Sahasrabudhe and Romario Thomas for their expert technical advice and assistance, and Dr. Almudena Bosch and Dr. Bridget Matikainen-Ankney for RNAseq data analysis. Microscopy was performed at the Microscopy Core and Advanced Bioimaging Center at Mount Sinai with assistance from Dr. Nikos Tzavaras and Glenn Doherty for FRAP imaging experiments and from Dr. Shilpa Dilip Kumar and Dr. Katarzyna Cialowicz for their guidance with super-resolution STED imaging, which utilized a microscope supported with funding from NIH Shared Instrumentation Grant (FAIN: S10OD021838). RNA extraction was performed by the Biorepository and Pathology Core at Mount Sinai and RNAseq was performed at Weill-Cornell Epigenomics Core.

## Notes

**Conflict of Interest:** The authors declare no competing financial conflicts.

### Competing Interest Statement

The authors have declared no competing interest.

