## Supplementary Figures & References for "Parkinson’s-linked LRRK2-G2019S derails AMPAR trafficking, mobility and composition in striatum with cell-type and subunit specificity"

A

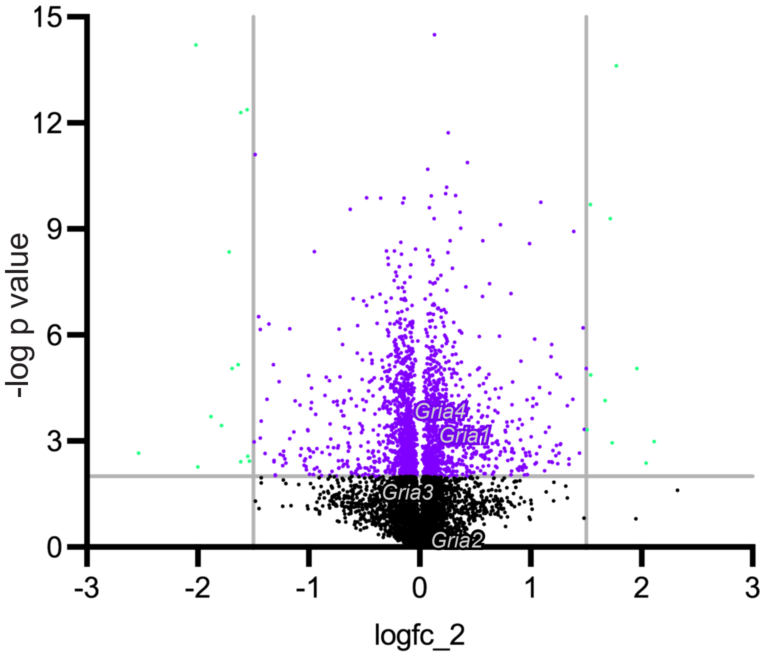


B

AMPAR Expression

| *Study (Bulk RNAseq)* | *Receptor mRNA* | *Log Fold Change* | *p value* | *q value* |
|---|---|---|---|---|
| *Lrrk2*$^{G2019S+/+}$ mouse striatum vs. *Lrrk2*$^{G2019S-/-}$ (wildtype) control (n=3 ea), 21 div (see methods for details) | Gria1 | 0.064 | 0.114 | 0.813 |
| | Gria2 | 0.010 | 0.888 | 0.981 |
| | Gria3 | -0.445 | 0.364 | 0.871 |
| | Gria4 | -0.124 | 0.080 | 0.796 |
| GSE183499<br>*LRRK2*$^{G2019S}$ iPSC patient derived differentiated toward neural stem cell vs. corrected control | Gria1 | -0.123 | 0.751 | 1 |
| | Gria2 | 0.844 | 0.299 | 1 |
| | Gria3 | -1.384 | 0.0980 | 1 |
| | Gria4 | 1.969 | 0.0006 | 0.333 |
| GSE136666<br>Parkinson Putamen vs.<br>nonPD control (3 Con, 3 PD) | Gria1 | -0.164 | 0.616 | 1 |
| | Gria2 | 0.035 | 0.017 | 1 |
| | Gria3 | 0.109 | 0.696 | 1 |
| | Gria4 | -0.495 | 0.155 | 1 |

**Supplementary Figure 1**. *LRRK2 mutation does not regulate AMPAR transcription*. Related to Figure 1. (**A**) Volcano plot shows the distribution of mRNAs sequenced from *wildtype (WT)* and *Lrrk2*$^{G2019S}$ *(GS)* striata (n = 3 males each; postnatal day 21). Lines on x-axis are set to mark < 1.5-fold change cutoff (green dots) and y-axis values are set to mark < 2 for statistical significance (purple and green dots). Few mRNAs (green dots) are significantly increased or decreased; and when p-values were corrected for false discovery rates (q value), only 2 candidates remained. Dots corresponding to mRNAs encoding AMPA receptors (Gria1 - 4) are annotated in gray. (**B**) Table compares AMPAR expression obtained from RNA sequencing experiments: *WT* and *GS* striata (outlined in A) and from prior work comparing iPSCs from *LRRK2*$^{G2019S}$ patient-derived stems cells and a corrected control (GEO DataSet GSE183499; Park S, Lee S, Chung S, 2021) and from RNA extracted from Parkinson's and control putamen samples (3 samples each; GEO DataSet GSE136666) (1) using GREIN (2). Across mouse and human tissue samples, when corrected for FDRs (q values) there were no significant differences in levels of mRNAs encoding AMPAR subunits.

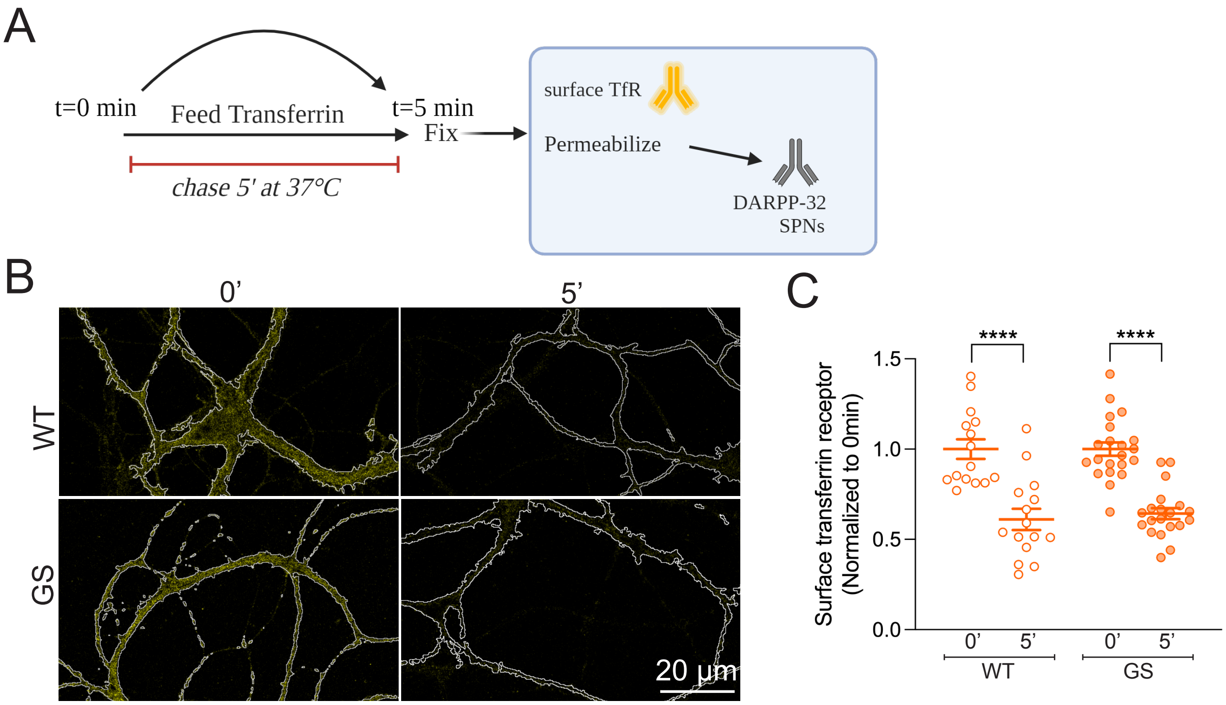


**Supplementary Figure 2**. *Clathrin-mediated endocytosis is intact in Lrrk2$^{G2019S}$ SPNs*. Related to Figure 2. (A) Outline of experiment in which surface TfR levels are evaluated at t = 0 mins and at t = 5 mins, following exposure to transferrin (30μg/ml) based on work showing that transferrin-bound transferrin receptors are rapidly endocytosed ($T_{1/2}$ = 3.5min) and recycled to the surface by ~15min (3). (**B**) Confocal images show surface labeling of TfR (yellow) within a mask generated using a threshold defined by DARPP-32 labeling to identify SPNs (and illustrated by white tracing). Magnification shown at lower right. (**C**) Scatterplot of surface TfR-labeling intensity normalized to t = 0 mins. Bars show ± SEM. 2-way ANOVA showed an effect of time, ****$p < 0.0001$; Holm Šidák post-hoc comparisons shown in graph; there was no effect of genotype or interaction. n= 15 - 21 neurons in 2 to 3 independent experiments.

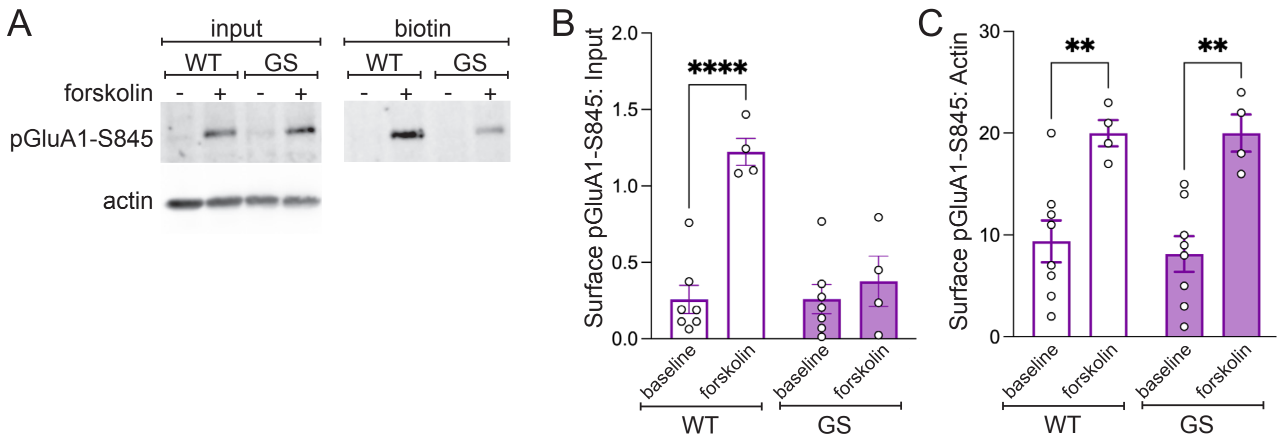


**Supplemental Figure 3.** *PKA mediated phosphorylation of GluA1 is intact in Lrrk2$^{G2019S}$ striatum***.** Related to data presented in Figure 3. **A**) Images of Western blots of input (left) and biotinylated (right) fractions of lysates generated from dorsomedial striatum dissections of wildtype (WT) and *Lrrk2*$^{G2019S}$ (GS) mice. Antibodies used (anti-pGluA1-S845 and actin) are indicated at left; and treatment conditions, baseline (-) and forskolin (+; 10min 50μM) are shown at top. **B**) Bar graph/ scatterplot shows quantification of westerns from biotinylated fractions. The biotin-accessible (surface) pGluA1-S845 protein levels expressed were normalized to input. Bars represent the mean ± SEM (n = 4-8 mice/genotype, 3 slices/mouse). Two-way ANOVA (F(1, 20) = 20.14, p=0.0002), *post hoc* Šidák's test ***p=0.0004 for WT. **C**) Bar graph/scatterplot shows pGluA1-S845 levels in the input fractions normalized to actin. Data were rank-transformed then analyzed by two-way ANOVA (F(1, 20) = 28.80, p<0.0001), *post hoc* Šidák's test, for WT **p= 0.0037, GS **p=0.0014.

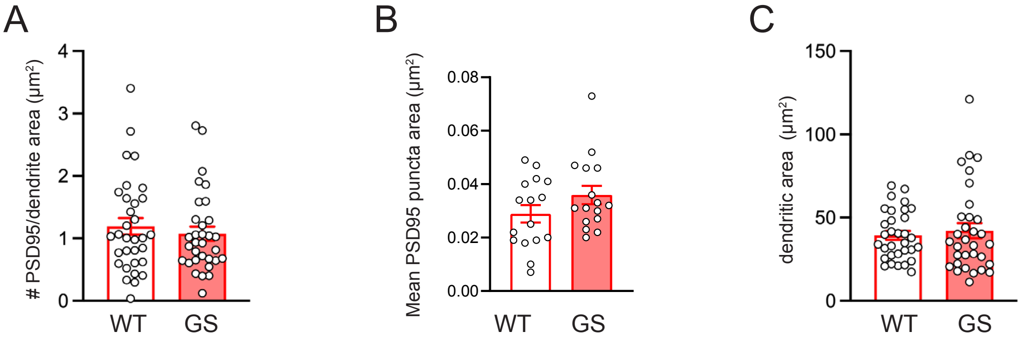


**Supplemental Figure 4.** *PSD95 density and area unchanged in Lrrk2*$^{G2019S}$ *$D_1$R-SPNs.* Bar graph/scatterplots compare wildtype (WT) and *Lrrk2*$^{G2019S}$ (GS) data generated from images captured using STED and confocal microscopy (as described in the Methods and in relation to Figure 4) using ImageJ. **A**) Mean density of PSD95 labeled puncta is similar across genotypes in $D_1$R-SPNs cultured for 18-21 days. (Mann Whitney test, p=0.6550). Data in (**B**) compare mean PSD95 puncta area within $D_1$R-SPN dendrites (unpaired t test, p=0.1475), and data in (**C**) compare mean tdTomato ($D_1$R-SPN) area sampled (Mann Whitney test, p=0.7439). Error bars are ± SEM.

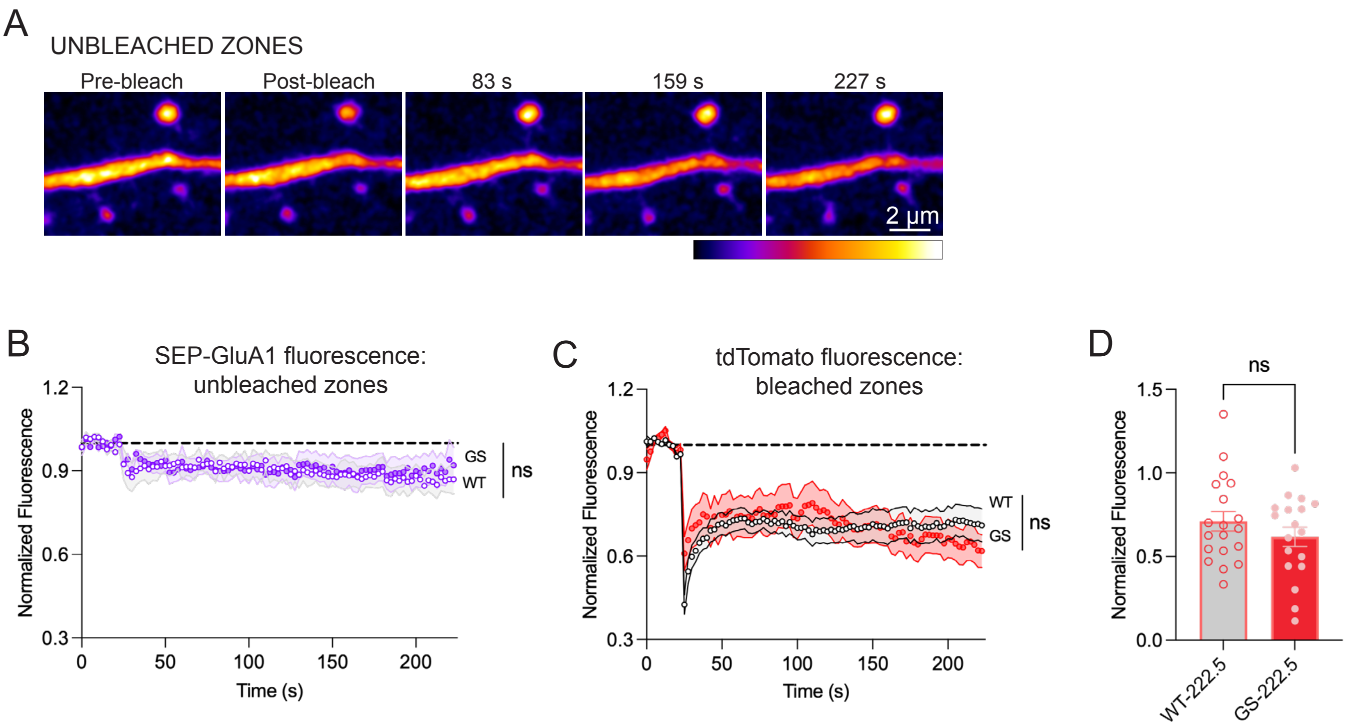


**Supplemental Figure 5.** *FRAP Controls.* Associated with Figure 4. (**A**) Confocal images of dendritic segments and SEP-GluA1 expressing spines that were not bleached, but imaged at the same intervals as data in Figure 5H to control for photobleaching. Look-up table used is shown beneath the images. (**B**) Plot shows no genotype-dependent difference (and little impact of photobleaching) in SEP-GluA1 fluorescence over time in unbleached ROIs. Wildtype (WT), open circles and *Lrrk2*$^{G2019S}$ (GS), filled purple circles. Shaded areas show ± SEM. Two-way ANOVA, impact of genotype: $p = 0.84$, $F(1,9) = 0.04$. (**C**). Plot of tdTomato (cytosplasmic) fluorescence recovery over time within ROIs that were bleached. There were no genotype dependent differences. WT, open circles and GS, filled, red circles; two-way ANOVA, impact of genotype: $p = 0.9$, $F(1,35) = 0.02$. (**D**) Bar/scatterplots compare degree to which fluorescence recovered at the end of the recording period (222.5 seconds). Error bars show ± SEM. Unpaired t test, $p = 0.28$.
